# Genetic diversity on the human X chromosome does not support a strict pseudoautosomal boundary

**DOI:** 10.1101/033936

**Authors:** Daniel J. Cotter, Sarah M. Brotman, Melissa A. Wilson Sayres

## Abstract

Unlike the autosomes, recombination between the X chromosome and Y chromosome often thought to be constrained to two small pseudoautosomal regions (PARs) at the tips of each sex chromosome. The PAR1 spans the first 2.7 Mb of the proximal arm of the human sex chromosomes, while the much smaller PAR2 encompasses the distal 320 kb of the long arm of each sex chromosome. In addition to the PAR1 and PAR2, there is a human-specific X-transposed region that was duplicated from the X to the Y. The X-transposed region is often not excluded from X-specific analyses, unlike the PARs, because it is not thought to routinely recombine. Genetic diversity is expected to be higher in recombining regions than in non-recombining regions because recombination reduces the effect of linked selection. In this study, we investigate patterns of genetic diversity in noncoding regions across the entire X chromosome of a global sample of 26 unrelated genetic females. We observe that genetic diversity in the PAR1 is significantly greater than the non-recombining regions (nonPARs). However, rather than an abrupt drop in diversity at the pseudoautosomal boundary, there is a gradual reduction in diversity from the recombining through the non-recombining region, suggesting that recombination between the human sex chromosomes spans across the currently defined pseudoautosomal boundary. In contrast, diversity in the PAR2 is not significantly elevated compared to the nonPAR, suggesting that recombination is not obligatory in the PAR2. Finally, diversity in the X-transposed region is higher than the surrounding nonPAR regions, providing evidence that recombination may occur with some frequency between the X and Y in the XTR.

## Introduction

The human sex chromosomes, X and Y, were previously an indistinguishable pair of autosomes, but within the last 180–210 million years, the ancestral pair diverged into two distinct chromosomes of tremendously different gene content and function^1,2^ The human sex chromosomes are composed of an older X-conserved region, shared across all therian (marsupial and eutherian) mammals ^3,4^ and a younger X- and Y-added region: autosomal sequence that was translocated to the X and Y chromosomes in the common ancestor of eutherian mammals approximately 80–130 million years ago ^5^. The differentiation of the X and Y is hypothesized to have occurred after a series of Y-specific inversions that suppressed X-Y recombination ^6–10^. In the absence of homologous recombination, the Y chromosome has lost nearly 90% of the genes that were on the ancestral sex chromosomes ^11–13^. Today the human X chromosome and Y chromosome share two pseudoautosomal regions (PAR1 and PAR2) at the ends of the chromosomes that continue to undergo homologous X-Y recombination ^6^. The PAR1 spans the first 2.7 Mb of the proximal arm of the human sex chromosomes ^11^ and contains genes from the ancient X- and Y-added region translocation. The PAR1 is separated from the non-recombining (nonPAR) region on the Y chromosome by a Y-specific inversion that is hypothesized to suppress X-Y recombination at this pseudoautosomal boundary ^10^. A functional copy of the XG gene spans the human pseudoautosomal boundary on the X chromosome ^14^, but is interrupted on the Y chromosome by a Y-specific inversion ^15^. In contrast to this mechanism for PAR formation, the 320kb human-specific PAR2 resulted from at least two duplications from the X chromosome to the terminal end of the Y chromosome ^16^.

Genes located in the PAR1 have important functions in all humans. Although genes on one X chromosome in 46,XX individuals are silenced via a process called X-inactivation ^17^ which evolved in response to loss of homologous gene content on the Y chromosome ^13^, all 24 genes in the PAR1 escape inactivation ^11,18,19^ (Supplementary Table 1). For example, one gene in the PAR1, SHOX1, plays an important role in long bone growth and skeletal formation ^20–22^. Consequence of SHOX1 disruption include short stature, skeletal deformities, Leri-Weill syndrome, and phenotypes associated with Turner syndrome (45,X) ^20^. ASMT, another gene located in the PAR1, is involved in the synthesis of melatonin, and is thought to be connected with psychiatric disorders, including bipolar affective disorder ^23^.

The suggested function of the pseudoautosomal regions is to assist in chromosome pairing and segregation ^24^. It has been proposed, in humans and great apes, that crossover events are mandatory during male meiosis ^25–27^ The analyses of human sperm suggest that a deficiency in recombination in PAR1 is significantly correlated with the occurrence of nondisjunction and results in Klinefelter syndrome (47, XXY) ^28^. Deletions in the PAR1 are shown to lead to short stature, which is correlated with Turner Syndrome ^29^. Further, the male sex-determining gene on the Y chromosome (SRY) is proximal to the PAR1 on the short arm of the Y. SRY can be translocated from the Y to the X during incongruent crossover events between the paternal PAR1s, resulting in SRY-positive XX males ^30^, or, more rarely, true hermaphroditism ^31^. The chances that XX individuals will inherit a copy of the SRY gene during male meiosis are restricted by reduced recombination at the PAR1 boundary ^32^.

Previous studies estimate that the recombination rate is approximately twenty times the genome average in the PAR1 ^26^ and about five times the genome average in the PAR2 ^33^, likely because recombination events in XY individuals are restricted to the pseudoautosomal sequences, with the exception of possible gene conversion in regions outside of the PARs ^34^ In addition to the PAR1 and PAR2, where recombination is known to occur between the X and Y, there is an X-transposed region (XTR) that was duplicated from the X to the Y in humans after human-chimpanzee divergence ^11,12^. Already, the XTR has incurred several deletions, and an inversion, but maintains 98.78% homology between the X and Y ^11^, and retains two genes with functional X-linked and Y-linked homologs ^12^.

Genetic diversity is expected to be higher in the pseudoautosomal regions than the remainder of the sex chromosomes for several reasons. First, recombination can unlink alleles affected by selection from nearby sites, reducing the effects of background selection and genetic hitchhiking on reducing genetic diversity ^35,36,62^. Second, the effective size of the pseudoautosomal regions of the sex chromosomes should be larger (existing in two copies in all individuals) than the non-recombining region of the X chromosome, which exists in two copies in genetic females and only one copy in genetic males. Finally, genetic diversity may be higher in pseudoautosomal regions than in regions that do not recombine in both sexes if recombination increases the local mutation rate^37–39^.

Studies of human population genetic variation often compare diversity on the X chromosome with diversity on the autosomes to make inferences about sex-biased human demographic history^40,41,42^. Typically the PAR1 and PAR2 are filtered out of these studies, at the defined pseudoautosomal boundaries, and the XTR is not filtered out. However, patterns of diversity across the entire human X chromosome, including transitions across the PARs and XTR have not been investigated to justify these common practices. In this study we investigate patterns of genetic diversity and divergence across the entire human X chromosome including the pseudoautosomal regions of twenty-six unrelated (46, XX) individuals sequenced to an average depth of 80X coverage by CompleteGenomics ^44^

## Results and Discussion

### Human X-linked nucleotide diversity is high in the PAR1 but not PAR2

We observe that uncorrected diversity is three times higher in the PAR1 than in the nonPAR regions, while uncorrected diversity in the PAR2 is not significantly greater than that in the nonPAR (Table 1; Figure 1; Supplementary Figure 1). We studied noncoding regions across the entire X chromosome, filtering out annotated genes, to minimize the effect of selection, but due to their small sizes, could not filter out regions far from genes in the PARs or XTR (Methods). However, mutation rate variation across the X chromosome may account for variable levels of diversity observed in the PAR and nonPAR regions. To correct for different mutation rates across the X chromosome, we normalize each region by divergence computed in that same region, with regions of high divergence filtered out, computed pairwise between human-chimpanzee, human-macaque, human-dog, or human-mouse (Supplementary Figure 2). Similar to the uncorrected diversity values, when we correct for mutation rate using any of these divergence values, we observe higher nucleotide diversity across humans in the PAR1 and PAR2 regions relative to the nonPAR regions, with diversity being significantly higher in the PAR1 than nonPAR (with XTR removed), and not significantly different between the PAR2 and nonPAR (Figure 1; Table 1; Supplementary Figure 1).

**Figure 1:**
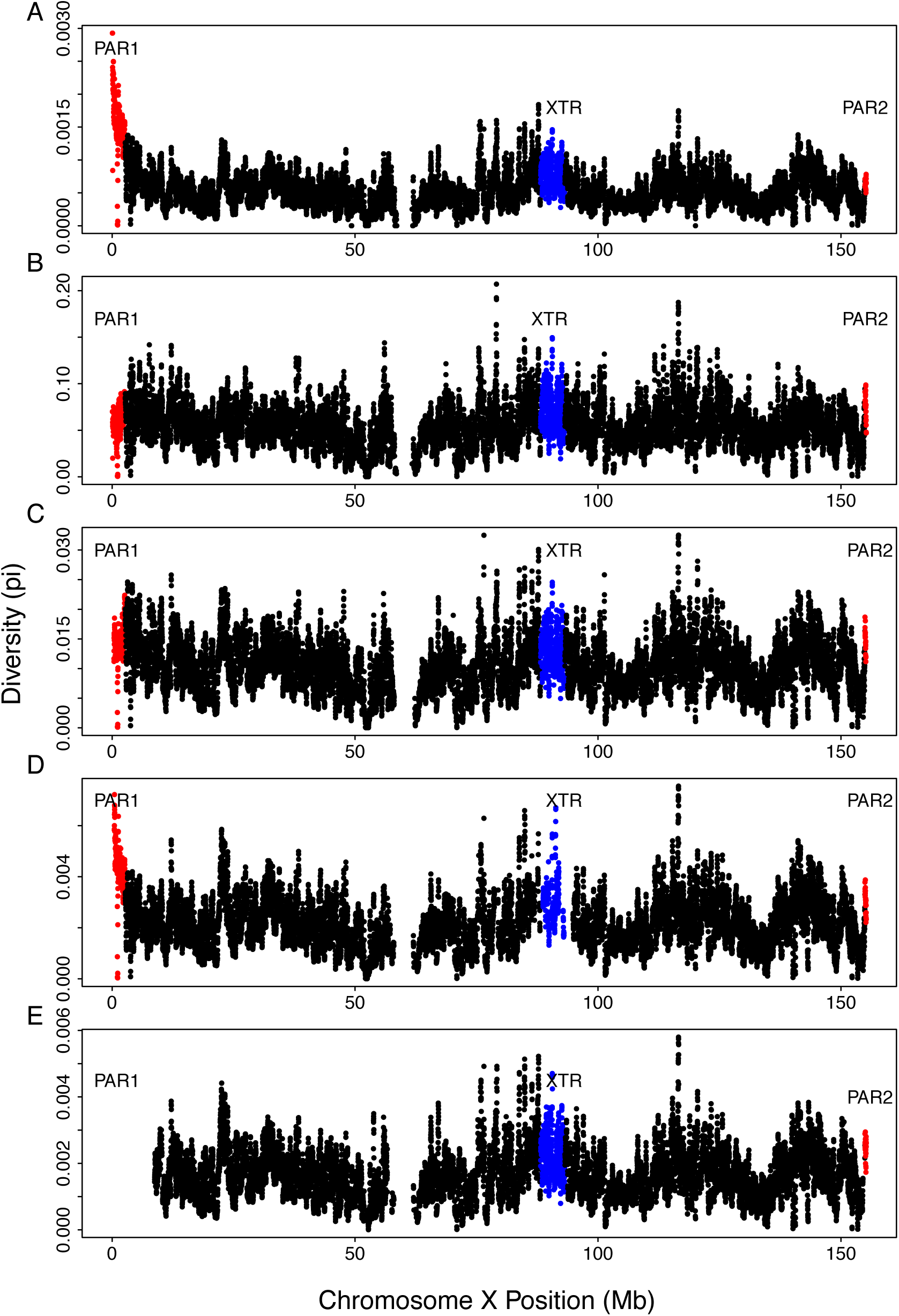
Diversity along the human X chromosome. Genetic diversity (measured by π) is shown in 100kb overlapping windows across the human X chromosome which includes the pseudoautosomal region 1 (PAR1), shown in red, the non-pseudoautosomal region, shown in black, the X-transposed region (XTR), shown in blue, and the pseudoautosomal region 2 (PAR2), shown in red, for A) human diversity uncorrected for divergence; then human diversity corrected for variable mutation rate using: B) human-chimpanzee divergence; C) human-macaque divergence; D) human-dog divergence; and E) human-mouse divergence.

**Table 1.**
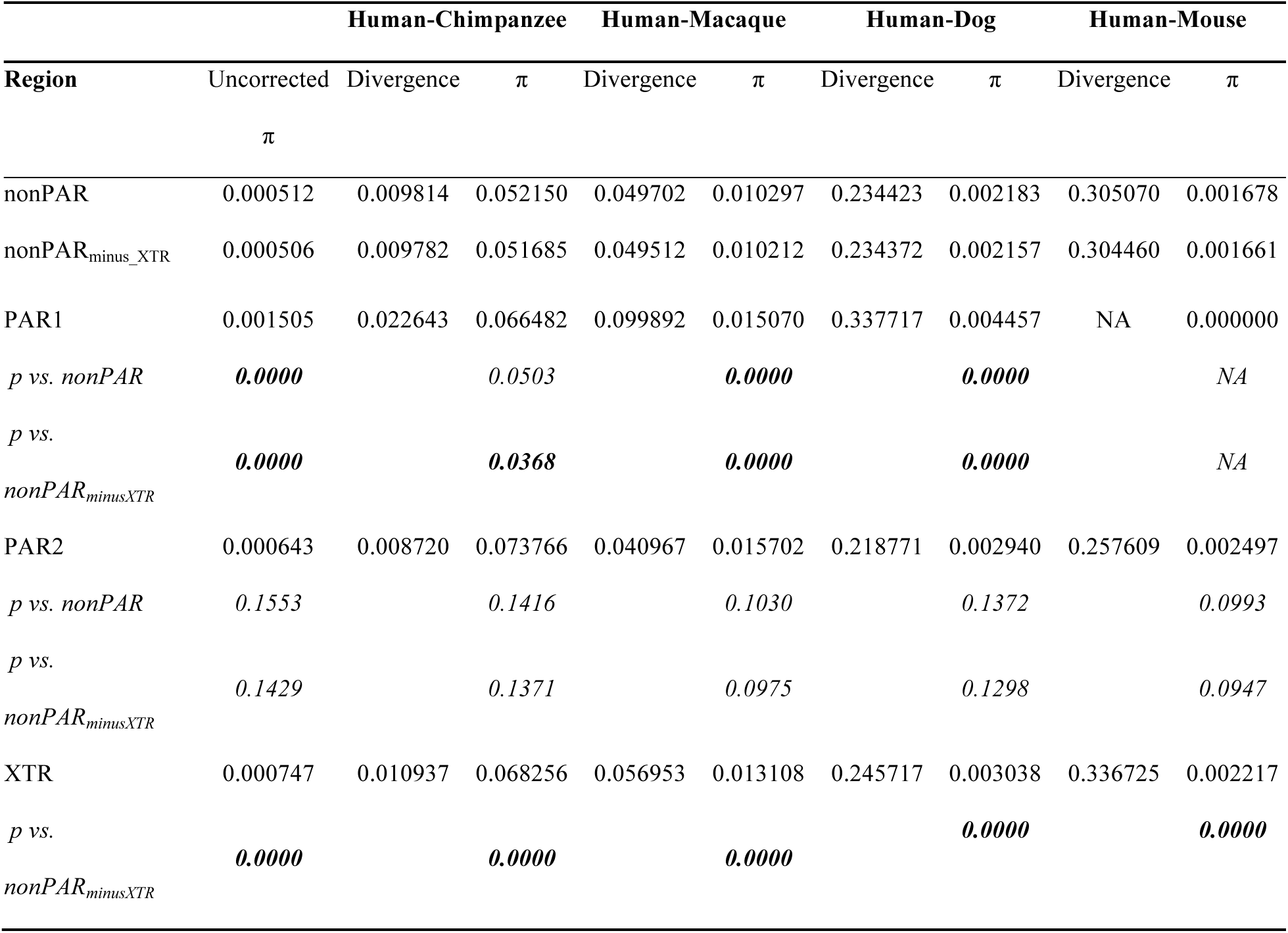
Diversity across regions of the human X chromosome. Diversity, measured as the average number of pairwise differences per site (π) between the X chromosomes of 26 unrelated genetic females, in each region of the human X chromosome is presented first unnormalized for mutation rate variation, then normalized using human-chimpanzee (hg19-panTro4) divergence and separately normalized for human-macaque (hg19-rheMac3) divergence. The regions analyzed include the pseudoautosomal region 1 (PAR1), pseudoautosomal region 2 (PAR2), X-transposed region (XTR) and the non-pseudoautosomal region either including the XTR(nonPAR) or excluding the XTR (nonPAR_minus___XTR_). *P* values from permutation tests with 1000 replicates are shown between each recombining and nonPAR region.

Curiously, human-chimpanzee and human-macaque divergence are quite high in the PAR1 relative to the nonPAR in a pattern that does not reflect diversity (Figure 1; Table 1). This results, predominantly, due to high inter-species divergence in the PAR1, and near the pseudoautosomal boundary (Supplementary Figure 3; Supplementary Figure 4). However, human-dog divergence roughly parallels uncorrected human diversity (Figure 1). Alignments between the human and mouse in PAR1 are unavailable.

Further, significantly elevated diversity in the PAR1 relative to the nonPAR cannot be attributed solely to mutation rate variation across the X chromosome, as the pattern remains after correction for divergence in each region (Figure 1; Table 1; Supplementary Figure 1). The pattern we observe is consistent with several processes, including: selection reducing variation more at linked sites in the nonPAR region than in the PAR1 due to reduced rates of recombination in the nonPAR relative to the PAR, or as a result of stronger drift in the nonPAR due to a smaller effective population size.

That we do not observe significantly elevated diversity in the PAR2 relative to the nonPAR is consistent with reports that the PAR2 undergoes X-Y recombination less frequently than the PAR1 ^48^, and supports assertions that in humans only one chiasma per chromosome is needed for proper segregation, rather than one per chromosome arm ^49^.

### Diversity is significantly higher in the X-transposed region than the non-pseudoautosomal region

Curiously, in addition to elevated rates of diversity in the previously described PAR1 and PAR2, we also observe that diversity is significantly higher in the recent X-transposed region, XTR, than in the nonPAR (Table 1; Supplementary Figure 1). This increased diversity cannot be attributed to mis-mapping between the X and Y as we only analyzed individuals with two X chromosomes (Methods). High diversity in the XTR contrasts with initial suggestions that there is no X-Y recombination in the XTR ^12^, and is consistent with recent reports of X-Y recombination in some human populations in this region ^46^.

Given the large size of the nonPAR region, and the small size of the XTR, 5Mb^11^, one may wonder whether removing the XTR would make a difference to measured levels of diversity across the human X chromosome. The raw diversity of the nonPAR including the XTR, measured as π, is 0.000512 while the raw diversity of the nonPAR excluding the XTR is 0.000506 (Table 1). Removal of the XTR does decrease both estimates of diversity and divergence in the nonPAR, and its inclusion in studies of X-specific diversity will affect inferences made when comparing X-linked and autosomal variation ^50–53^.

### Pseudoautosomal boundaries cannot be inferred from patterns of diversity

Recombination between the X and Y is expected to be suppressed at the pseudoautosomal boundary, where X-Y sequence homology diverges due to a Y-specific inversion ^10,14,15^. If diversity correlates highly with recombination rate, and X-Y recombination is strictly suppressed in the nonPAR after the pseudoautosomal boundary, then diversity is expected to drop sharply between the PAR1 and nonPAR. However, when we analyze patterns of human diversity in windows across the X chromosome (Methods), we do not observe an abrupt shift in the level of diversity between the PAR1 and nonPAR regions (Figure 1). The lack of an observable pseudoautosomal boundary based on diversity is clear whether small or large (100kb or 1Mb), or overlapping or non-overalapping windows are used (Methods; Supplementary Figure 7). The history of gene conversion between the sex chromosomes may contribute to the increased diversity levels^54^ in the nonPAR side of the Y-specific inversion that marks the pseudoautosomal boundary. Human diversity uncorrected for divergence decreases from the proximal end of the PAR1 through the pseudoautosomal boundary and well into the nonPAR. A sex-specific map of the PAR1 found that male recombination is higher near the telomeres, and decreases near the pseudoautosomal boundary, while, in contrast, the female recombination rate reported in the same study in the PAR1 is fairly flat throughout the region and increases near the pseudoautosomal boundary ^55^. Thus, genetic diversity uncorrected for divergence in the PAR1 appears to correlate with the male recombination rate. Curiously, however, a previous study of recombination rate in the PAR1 reported an increase in the female (but not the male) recombination rate near the proximal end of the PAR1 ^56^. Thus, potentially both male and female recombination rates contribute to the linear decrease in diversity observed in the PAR1 from the proximal end of the X through the pseudoautosomal boundary. Although not yet mapped, when the data becomes available, it will be useful to compare patterns of diversity with sex-specific recombination maps across the entire X chromosome.

## Conclusions

We show that diversity is indeed higher in the pseudoautosomal regions, and lower in the regions of the X that are not known to recombine in males (nonPAR). Diversity in the PAR1 is significantly higher than in the nonPAR regardless of normalizing the diversity with divergence between human and either chimpanzee, macaque, or dog to correct for mutation rate (Table 1; Figure 1). Diversity was also normalized with divergence from the mouse, but there is not alignment between human and mouse in the PAR1, due to a different evolutionary origin in the PAR1 and no common pseudoautosomal genes are shared between them ^45^. We observe that diversity is lower in the PAR2 than expected, and is not significantly different from the nonPAR region. We also show that diversity is elevated in the XTR above other nonPAR regions, verifying recent observations that the region may still undergo homologous recombination between the X and Y chromosomes ^46^. Finally, when analyzing patterns of genetic diversity in windows across the human X chromosome we find that there is no strict boundary, based solely on the levels of diversity, between the recombining and putatively non-recombining regions, which could be attributed to the evolutionary shift in the PAB over time, extending the PAR1 due to a PAR1 length polymorphism ^47^ This could also suggest that non-homologous recombination at the pseudoautosomal boundaries may be common.

Our observations of patterns of diversity across regions of the human X chromosome with variable levels of recombination are consistent with previous reports that diversity and divergence are correlated with recombination rate in humans across the genome ^38^, and explicitly in the PAR1 ^57^ Elevated levels of diversity in the XTR suggest that, consistent with a recent report ^46^, this region may frequently undergo X-Y recombination. Curiously, we did not find a significant elevation in diversity in the PAR2, that, in agreement with its unusual evolution^16^, indicate that it rarely recombines between X and Y during meiosis. Further, the lack of a clear differentiation in diversity between the PAR1 and nonPAR suggests that recombination suppression between X and Y is still an actively evolving process in humans, as in other species^58^. This is consistent with evidence that the position of the pseudoautosomal boundary varies across mammals ^59–62^. There is even evidence of polymorphism in the pseudoautosomal boundary in a pedigree analysis of a paternally inherited X chromosome in humans ^47^. Taken together with previous inferences about the variation in pseudoautosomal boundaries, our observations suggest that assumptions should not be made of a strict suppression of X-Y recombination at the proposed human pseudoautosomal boundary.

## Methods

We analyzed X chromosomes from twenty-six unrelated (46, XX) individuals sequenced by CompleteGenomics ^44^ (Supplementary Table 4). Sites were filtered requiring data be present (monomorphic or variable) in all twenty-six samples. Human-chimpanzee (hg19-panTro4), human-macaque (hg19-rheMac3), human-dog (hg19-canFam3), and human-mouse (hg19-mm10) alignments were extracted from the UCSC genome browser ^63^. We curated the human-chimpanzee and human-macaque alignments to filter out segments that included autosomal sequence aligning to the X chromosome (Supplementary Table 2; Supplementary Figure 5; Supplementary Figure 6); These alignments were visualized using the gmaj software ^64^ Additionally, we observed several regions across the X chromosome exhibiting heightened divergence between the human and the chimpanzee or the human and the macaque (Supplementary Figure 3; Supplementary Figure 4). Upon further inspection, these regions often contain multi-copy gene families that could lead to mismapping (Supplementary Table 2).

Divergence estimates were similar with and without these regions (Supplementary Table 3), and here we present results with these regions of high divergence near multi-copy gene families excluded. Some regions of the X chromosome exhibit extremely high inter-species divergence (Supplementary Figure 3; Supplementary Figure 4; Supplementary Table 2). These regions were excluded from this analysis, but we reach the same conclusions described above whether they are included or excluded.

We used the Galaxy tools ^65^ to filter out regions that could cause potential sequence misalignments and regions defined by the UCSC genome browser ^63^ that may be subject to selection: RefSeq genes, simple repeats, and repetitive elements. We attempted to filter out noncoding regions near genes, but doing so would leave very little analyzable sequence in the PAR1 and PAR2 regions.

We measured the diversity between the sequences as π, the average pairwise nucleotide differences per site between all sequences in the sample.

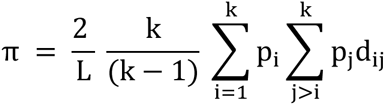

L represents the number of called sites, *k* represents the number of DNA sequences, *p_i_* and *p_j_* are the frequencies of the corresponding alleles *i* and *j*, and *d_j_* is the number of sites containing nucleotide differences. Diversity was calculated within each specific region (PAR1, PAR2, XTR, nonPAR with XTR, and nonPAR without XTR), as well as across sliding and non-overlapping windows. We generated window interval files across the human X chromosome with Galaxy tools ^65^, and conducted analysis, in four sets of windows (Supplementary Figure 7): 1) in a 1 Mb non-overlapping window, 2) a 1 Mb window with 100 kb sliding start positions, 3) a 100 kb non-overlapping window, and 4) a 100 kb window with 10 kb sliding start positions. We similarly calculated human-chimpanzee, human-macaque, human-dog, and human-mouse species divergence along the X chromosome, in each of the four regions and in the same windows previously described. To normalize the data, π values were divided by the observed divergence within the same interval.

Chromosome X was divided into windows that were permutated without replacement 10,000 times to assess significant differences between diversity in each region (PAR1, nonPAR, XTR, PAR2). This analysis was repeated for uncorrected diversity and diversity corrected for human-chimpanzee, human-macaque, human-dog, and human-mouse divergence values. Empirical *P-*values were calculated by comparing differences between permutated samples with differences in observed diversity between each pair of regions. The negative correlation along the pseudoautosomal boundary was tested using linear regressions across 100kb windows covering a total of 3Mb for each regression (30 windows), shifting the window by 100 kb systematically (Figure 2). Each regression was analyzed for significance (p < 0.05) with all data points occurring before the first non-significant window being included in the significant dataset. All of the graphs were produced using R version 3.1.2 ^66^. All codes used for this project can be found at: https://github.com/WilsonSayresLab/PARdiversity.

**Figure 2:**
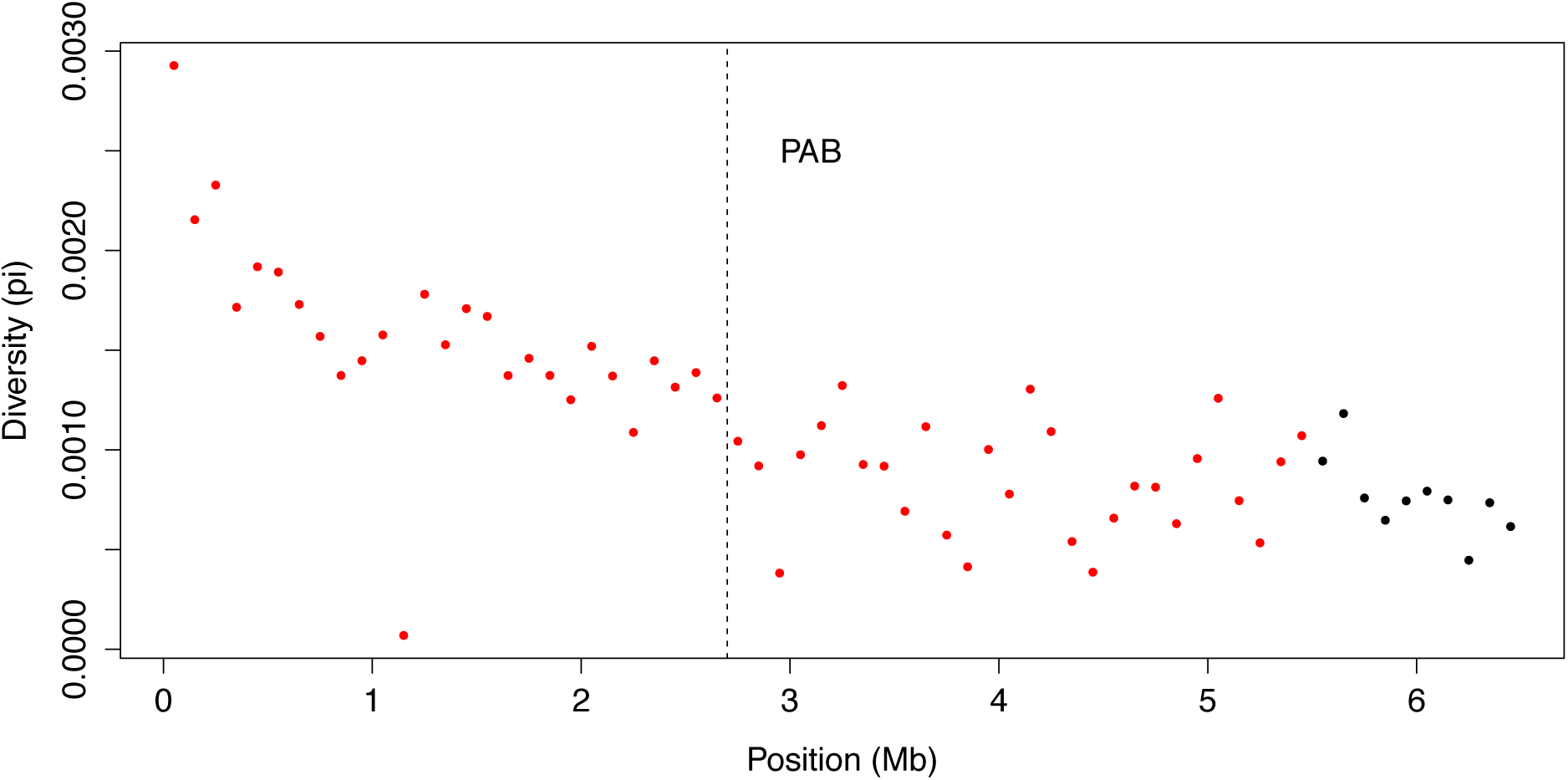
Negative correlation between diversity and distance from Xp, crossing the pseudoautosomal boundary. Diversity in 100kb non-overlapping windows along the psuedoautosomal boundary (PAB) is plotted across the first 6Mb of the human X chromosome, spanning the annotated pseudoautosomal boundary at 2.7Mb. A series of linear regressions were run including 30 windows, sliding by one window across the PAR to the nonPAR region. Each 100kb window is colored red if it is included in a regression in which distance from Xp and diversity are significantly negatively correlated, otherwise the windows are colored black. For the entire region together, diversity is significantly negatively correlated with distance from Xp (*p* = 3.281 × 10^−10^; r = −0.7321563) and spans the pseudoautosomal boundary.

**Figure 3:**
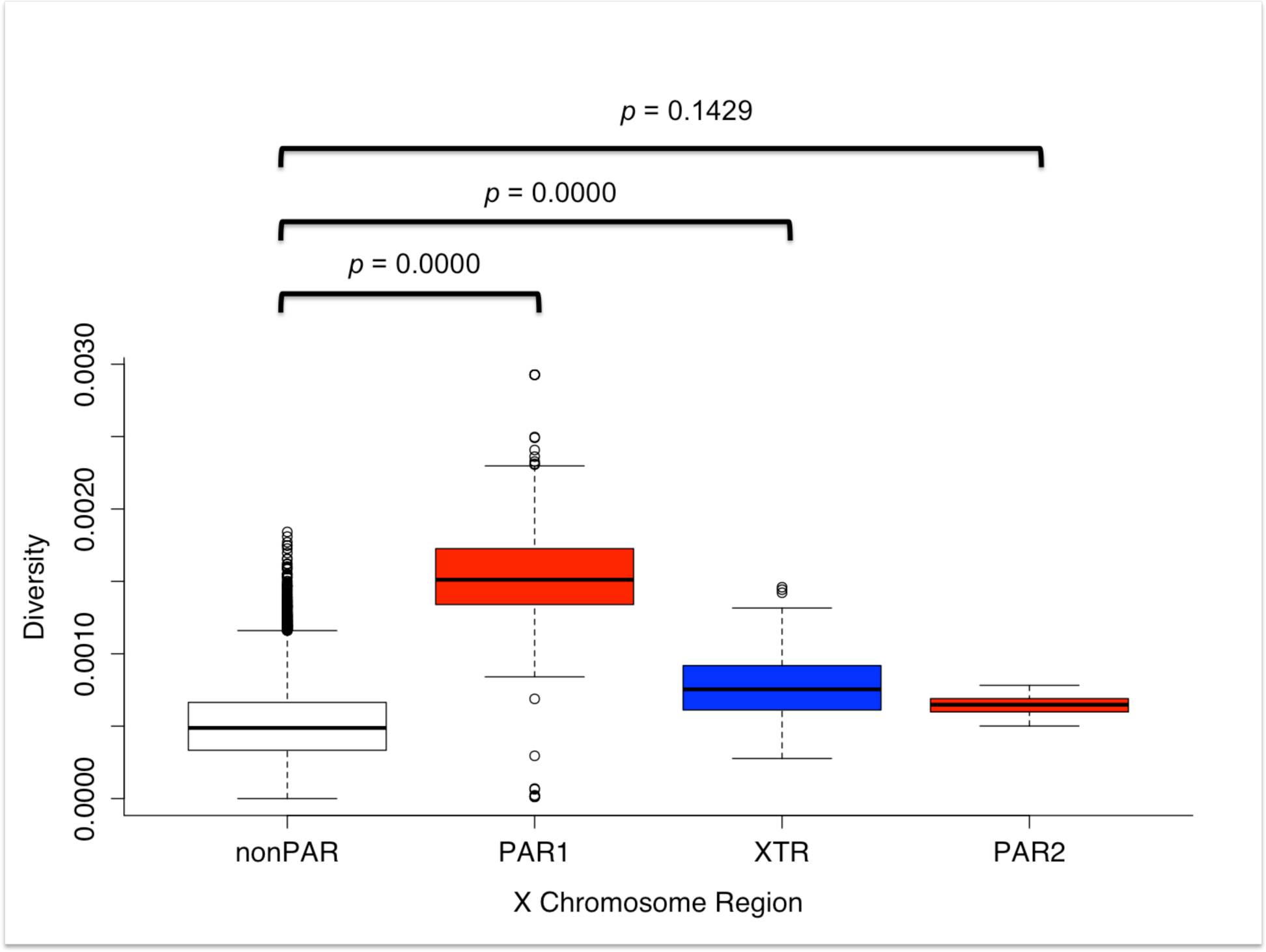
Diversity along the X chromosome split by region. Genetic diversity (measured by π) is shown in boxplots depicting the average diversity with error bars for the nonPAR, PAR1, XTR, and PAR2. The *p value* from a permutation test with 1000 replicates, comparing the diversity of each region to the diversity of the nonPAR are shown.

